# A general methodology for liver sinusoid fenestration analysis based on 3D electron microscopy data

**DOI:** 10.64898/2026.03.07.710307

**Authors:** Cécile Pohar, Yousr Rekik, Minh Son Phan, Benoit Gallet, Agnès Desroches-Castan, Mireille Chevallet, Jean-Yves Tinevez, Emmanuelle Tillet, Nicola Vigano, Pierre-Henri Jouneau, Aurélien Deniaud

## Abstract

The liver has a complex architecture composed of millions of lobules. Within these lobules, hepatocytes, the main hepatic cells, are organized in rows separated by blood capillaries known as sinusoids. These capillaries are lined by liver sinusoidal endothelial cells (LSEC) that form a very specific fenestrated endothelium essential for the exchange of metabolites and proteins between the blood and hepatocytes. Alterations in the size and number of LSEC fenestrations are associated with the onset and the progression of various liver diseases. The analysis of liver architecture is thus of utmost importance for advancing our knowledge of liver ultrastructure and its alterations.

Liver architecture has been studied since decades, mainly using 2D electron microscopy, and more recently using advanced super-resolution fluorescence microscopy. In recent years, volume electron microscopy techniques, including focused ion beam-scanning electron microscopy (FIB-SEM) progressed and nowadays enable the 3D reconstruction of biological ultrastructures down to nanometer resolution. However, the analysis of large volumes (e.g., several tens of µm^3^) remains challenging due to various constraints in the segmentation of large datasets. In the current study, we developed a workflow to semi-automatically segment hepatic sinusoids from FIB-SEM mice liver datasets using the CNN-based (convolutional neural network) tool known as “nnU-Net”, after fine-tuning a ground truth model. We also implemented tools for semi-automatic quantification of LSEC fenestrae diameters and sinusoid porosity from segmented datasets. This workflow enabled us to compare the distribution of LSEC fenestrae diameters in wild-type versus *Bmp9*-deleted mice, a hepatic factor known to be involved in fenestration maintenance. Our results confirm the importance of BMP9 for LSEC differentiation. Therefore, the developed methodology represents a valuable tool for characterizing the fenestrated endothelium under various physiological and pathological conditions.

## Introduction

The liver is a major organ in humans, performing many metabolic functions that are important both for nutrient storage and distribution and for xenobiotic transformations and excretion into the feces via the bile. This central role is made possible by its architecture and the specific polarity of hepatocytes, the main cell type of the liver, which represent around 80% of its mass [1]. Indeed, hepatocytes have three distinct types of poles, i) basal poles, contacting the sinusoids, ii) apical poles, forming the bile canaliculi, and iii) lateral poles, located between adjacent hepatocytes and separating the basal and apical poles. Sinusoids are small blood capillaries lined with Liver Sinusoidal Endothelial Cells (LSEC) [2], where blood rich in both oxygen and nutrients mixes before draining into the central vein, through which it is transported out of the liver. Nutrient-rich blood, which is crucial for the liver metabolic functions, arrives from the intestine *via* the portal vein. Sinusoids possess a highly specialized, porous endothelium that is in direct contact with the underlying hepatocyte microvilli, allowing an efficient nutrient uptake by hepatocytes. Indeed, LSEC are perforated with transmembrane pores, called fenestrations or fenestrae, whose diameters vary between 50 and 300 nm [3]. Fenestrations lack a diaphragm and allow the bidirectional exchange of small molecules and proteins, such as lipoproteins between blood and hepatocytes through the space of Disse, while filtering out larger molecules. This size-dependent transport makes LSEC an efficient ultrafiltration system.

In various hepatic diseases as well as upon aging, the sinusoidal endothelium undergoes defenestration and/or capillarization characterized by the loss of fenestrations and the deposition of a basal membrane. In chronic hepatic diseases, these alterations usually occur concomitantly with fibrosis, marked by excessive deposition of extracellular matrix in the space of Disse, leading to a decrease in LSEC porosity and thereof to reduced molecule exchanges between hepatocytes and the blood [4,5]. For instance, it has been shown that the deletion of *Gfd-2,* the gene coding for the circulating growth factor BMP9 produced by the hepatic stellate cells leads to a slight increase in the diameter of fenestrae and a three-fold decrease in their number in the mouse sinusoidal endothelium [6]. These results show the importance of BMP9 on LSEC differentiation and fenestration maintenance.

Atomic force microscopy experiments performed on isolated living mouse LSEC have shown that fenestrae are dynamic structures able to modulate their diameter within minutes to hours [7]. Super-resolution optical microscopy techniques have provided complementary insights into the ultrastructure of fenestrae and their spatial organization under physiological and pathological conditions, as well as following treatments with various agents (toxins, drugs…) that influence their diameter and/or number [8].

The most standard methods for the analysis of LSEC fenestrae are electron microscopy techniques performed on liver blocks, sections or isolated LSEC [9,10]. However, a three-dimensional analysis of hepatic sinusoids at nanometer resolution and in the hepatic context has not yet been achieved. Such an approach is essential to obtain accurate quantitative information on the number, diameter, and spatial distribution of LSEC fenestrae, parameters of utmost importance for understanding sinusoidal function in health and disease.

To achieve this goal, we developed a methodology based on volume electron microscopy by Focused Ion Beam-SEM (FIB-SEM), performed on mouse liver samples to acquire three-dimensional data at nanometer scale. To obtain information as close as possible to the native state, liver sample preparation was optimized, including a high-pressure freezing step, which increased the contrast between sinusoids and the underlying hepatocyte microvilli. This step was essential to segment the sinusoid in a multi-step workflow taking into account the structural specificities of LSEC, which include both thin fenestrated regions and large nuclear parts. Classical segmentation approaches such as thresholding struggle to handle the specific type of contrast found in EM images of hepatic sinusoids. Indeed, the limited difference in local contrast between the membranes of LSEC and the surrounding cellular structures prevents accurate delineation using direct methods. Traditional machine learning-based methods represent a rational alternative, as they can learn the relevant features defining the contours of the objects of interest. However, while helpful for annotations and manual refinement, these approaches lacked robustness when propagating segmentation across large volumes, leading to numerous false positives inside cells. Therefore, these methods, combined with dedicated post-processing steps, were used to generate a ground truth segmentation of a hepatic sinusoid. Extending this segmentation approach to other sinusoids required a more generalizable strategy, motivating the adoption of deep learning-based methods. Deep learning algorithms, in particular convolutional neural networks (CNN), have significantly evolved in recent years. The U-Net architecture has become the reference architecture for image segmentation tasks due to its versatility and ability to capture multiscale contextual information [11]. Among the available U-Net-based frameworks, nnU-Net was selected for the segmentation of LSEC thanks to its ability to automatically adapt its hyperparameters to the characteristics of the image dataset at hand [12].

Finally, we developed tools to semi-automatically quantify both the number and the diameter of LSEC fenestrae based on the sinusoid segmentation. Altogether, an integrated workflow covering the entire process from liver sample preparation to the extraction of quantitative information on sinusoid fenestrations was established. This workflow can be applied to various liver samples, as demonstrated with BMP9-deleted mice (name hereafter *Bmp9-KO* mice). This approach is therefore of high interest for comparing sinusoid fenestrations in different biological contexts.

## Methods

### Mouse liver sample preparation

All animal procedures were conducted in accordance with the institutional guidelines of the European Community (EU Directive 2010/63/EU) and approved by the CEA local ethics committee and the French Ministry of Higher Education and Research (APAFIS # #9436-2017032916298306). Mice were from a 129/Ola genetic background with a wild-type (WT) or *Bmp9*-KO genotypes as described in [6].

The mice were lethally anesthetized using pentobarbital injection (180 mg/kg) and then perfused through the left ventricle with PBS buffer followed by 10-15 mL of fixative buffer (2.5% glutaraldehyde and 1.5% paraformaldehyde in 0.1 M cacodylate buffer, pH 7.4, containing 2 mM calcium chloride CaCl₂). The liver was then harvested and cut into small cubes of approximately 5 × 5 × 5 mm³ that are kept in fixative until freezing.

For the optimized protocol, including cryo-fixation and cryo-substitution, the liver cubes were placed on a type A aluminum high-pressure freezing carrier of 3 or 6 mm in diameter (Leica Microsystems) in a few drops of cryo-protectant (20% BSA). This freezing cup was then covered on the flat side with another type B freezing cup of 3 or 6 mm in diameter (Leica Microsystems) and the assembly was introduced into an HPM100 cryofixator (Leica Microsystems). This method uses liquid nitrogen (T = −196°C) at high pressure (210 MPa) to fix the sample.

After cryo-fixation, the samples were transferred to an automated freeze substitution system (AFS2, Leica Microsystems). The freeze substitution cycle used was as follows. First, the samples were incubated in an acetone bath supplemented with 2% osmium tetroxide and 0.5% uranyl acetate in dried acetone at −90°C for 40 h. Then, they were slowly warmed to −60°C (1°C/h) and maintained at this temperature for 12 h. Next, the temperature was gradually increased to −30°C at a rate of 1°C/h. After 12 h, the temperature was rapidly raised to 0°C and the samples were maintained for 1 h at this temperature. Again, the samples were cooled to −30°C then rinsed four times in pure acetone for 15 minutes at the same temperature. The samples were then resin-embedded by incubation in three successive baths of increasing concentrations of araldite *Durcupan ACM Epoxy* resin (*Electron Microscopy Science*, EMS, 14040) in acetone. They were left for 2 h in each bath while gradually increasing the temperature from −30°C to 20°C: 2 h in a 1:2 (volume:volume) resin/acetone mixture while raising the temperature from −30°C to −10°C; 2 h in a 1:1 resin/acetone mixture while raising the temperature from −10°C to 10°C; and 2 h in a 2:1 resin/acetone mixture while raising the temperature from 10°C to 20°C.

The samples were then incubated in a pure resin bath at room temperature for 8-10 h, followed by a mixture of pure resin and the accelerator benzyldimethylamine for 8 h at room temperature. Finally, the resin was polymerized at 60°C for 48 h.

### FIB-SEM data acquisition

3D FIB-SEM stacks were acquired using a Zeiss crossbeam 550 microscope. After trimming with an ultramicrotome (Leica UC6) to obtain a flat surface, the resin block was mounted on an aluminium stub with silver paste, fully coated with a 5-nm-thick platinum layer for conductivity (Safematic CCU-010), and transferred to the microscope for acquisition with the Atlas 5 software (Zeiss). Two converging fiducial lines were engraved between a platinum layer and a carbon layer deposited on the target area. A large trench was then opened in front of this region using a 30 kV / 30 nA Ga^+^ beam. Image stacks were obtained by simultaneously milling the surface with the Ga^+^ beam and recording backscattered electron SEM images with the in-column EsB detector. Fiducial lines were used to dynamically compensate for drift and to monitor slice thickness accuracy.

FIB-SEM enables isotropic volume acquisition, with slice thickness matching the SEM pixel size. A trade-off must be made between resolution, total volume size, and acquisition time. In this study, samples were imaged at high resolution (4 nm × 4 nm × 4 nm per voxel). Typical acquisition parameters were 30 kV / 700 pA or 1.5 nA for Ga^+^ milling, and 1.5 kV / 1 or 1.5 nA / 5 - 10 µs dwell time for SEM acquisition with the EsB detector, with an EsB grid voltage of 600 V. Depending on the size of the area of interest, the dwell time is chosen so that the acquisition time per frame remains below 2 minutes for better stability.

### Preprocessing of FIB-SEM image stacks

FIB-SEM image stacks were aligned in Fiji [13] using MultiStackReg [14] or Scale Invariant Feature Transform (SIFT) [15] plugins. MultiStackReg aligns each image with respect to the previous one using cross-correlation and SIFT is based on the detection and matching of local image features between successive slices. For MultiStackReg, the first or the central image was used as a global reference for the whole stack and translation was used as the transformation type. After alignment, the image stack was checked in orthogonal views and if a misalignment persisted, the angle of misalignment was corrected using a home-made Python script.

### LSEC ground truth segmentation for WT mouse liver samples

The ground truth segmentation was generated from an image stack of a sinusoid acquired on a WT mouse liver block.

#### Step 1: Definition of the LSEC fenestration localization region

This step aimed to define the LSEC fenestrated region, corresponding to the area containing fenestrations and serving as a mask for subsequent segmentation steps.

The initial FIB-SEM image stack was first binned in Fiji to achieve a voxel size of 64 × 64 × 64 nm³. The binned image stack was then imported into ilastik (version 1.33 or 1.4b) [16] to segment the sinusoid lumen using the *Carving* workflow. The resulting segmentation mask was resampled in Fiji using bicubic interpolation to recover the original isotropic resolution of 4 nm in all three directions. Post-processing of this resampled mask was then performed in IPSDK Explorer (Reactiv’IP) in the following order:

- Binarization of the mask
- Application of the *outline* filter to extract the sinusoid contour
- Application of a 3D dilation using a spherical structuring element with a radius sufficient to encompass the LSEC fenestrations

Finally, a Gaussian filter was applied to generate the mask defining the fenestration localization region.

#### Step 2: Segmentation of the LSEC fenestrated regions

The initial image stack and the mask of the fenestration localization region (result of step 1) were combined in Fiji to form a 2-channel image stack. Two sub-volumes representing a fenestrated and a nuclear region of LSEC were then extracted and converted to HDF5 format before being imported into ilastik for segmentation using the *Pixel Classification* workflow. After segmentation of these two sub-volumes, the segmentation was generalized to the entire image stack. The resulting binary mask (segmentation mask 1) was post-processed in Fiji to refine the segmentation following these steps:

- Inversion and 3D dilation of the fenestration localization region mask to generate a subtraction mask.
- Subtraction of the subtraction mask from the segmentation mask 1.

The result of this step was the segmentation mask of the LSEC fenestrated regions.

#### Step 3: Segmentation of the LSEC nuclear regions

In this step, images were first preprocessed to define the LSEC localization region, followed by segmentation of the LSEC nuclear regions.

##### Initial image preprocessing

The LSEC localization region corresponds to the area occupied by LSEC around the sinusoid contour, defined in step 1. It was subdivided into two parts: the LSEC luminal region extending towards the sinusoidal lumen and the LSEC parenchymal region extending towards the hepatocytes.

###### Determination of the LSEC parenchymal region

In IPSDK, the segmentation mask of the sinusoid lumen (step 1) was dilated in 3D using a spherical structuring element sufficient to encompass the extension of LSEC towards hepatocytes. The non-dilated mask was then subtracted from the dilated one, defining the LSEC parenchymal region (**Figure S1**).

###### Determination of the LSEC luminal region

The following steps were performed in IPSDK (**Figure S2**):

- Dilation of the segmentation mask of the sinusoid lumen using a spherical structuring element with a radius sufficient to encompass the extension of LSEC (radius 1) towards the sinusoid lumen
- Subtraction of the non-dilated mask from the dilated mask. The result was called the intermediate mask.
- Dilation of the sinusoid contour, defined in step 1, using a spherical structuring element of radius 1.
- Subtraction of the intermediate mask from the dilated contour mask. The result defined the LSEC luminal region.

###### Determination of the LSEC localization region

A pixel-wise addition of the LSEC parenchymal and luminal regions was performed in Fiji to define the LSEC localization region (**Figure S3**).

###### Application of the LSEC localization region to the initial images

The following steps were performed in Fiji (**Figure S4**):

- Subtraction of the initial data from the LSEC localization region: LSEC localization region - initial data
- Contrast inversion of the subtraction result

Hepatocytes were masked in the preprocessed initial stack, leaving only LSEC for subsequent segmentation.

##### Segmentation of the LSEC nuclear regions

The preprocessed initial images were binned with a factor of 8 in all three directions, resulting in a voxel size of 32 × 32 × 32 nm³. The binned stack was imported into ilastik to segment LSEC nuclear parts using the *Carving* workflow.

The resulting segmentation mask was then resampled in Fiji using the cubic interpolation to restore the original isotropic resolution (4 nm × 4 nm × 4 nm per voxel). This mask defined the LSEC nuclear regions.

#### Step 4: Final LSEC segmentation

The preprocessed initial images (voxel size = 4 × 4 × 4 nm³), the LSEC fenestrated regions mask, and the LSEC nuclear regions mask were combined in Fiji to form a 3-channel image stack. In total, six sub-volumes were extracted from this image stack and converted to HDF5 format before being imported into ilastik via the *Pixel Classification* workflow.

After training the classifier on the sub-volumes, the segmentation was generalized to the entire image stack. The dimensions of the sub-volumes used in the training are detailed in **Table S1**.

##### LSEC ground truth segmentation for *Bmp9-KO* mice samples

The pipeline designed for WT mice livers was adapted to take into account the specificities of *Bmp9-KO mice* livers. The main difference lies in the number of classes used in Pixel Classification to generate the ground truth segmentation. Three classes, background, object of interest corresponding to the LSEC and everything else were defined in Pixel Classification for samples from *Bmp9-KO* mice, compared to only two (i.e., LSEC and everything else) for samples from WT mice, as described above.

##### Generation of the final mesh

The segmentation masks obtained after ilastik training on 4 and 6 sub-volumes followed by segmentation generalization to the entire image stack: sub-volumes 1–4 and 1–6 (**Table S1**) were exported, converted to TIFF, and combined in Fiji using an arithmetic OR operation. The resulting mask was imported into IPSDK, where a connected components analysis was applied, and the largest connected component was retained. A 3D mesh of this component was then generated. Mesh smoothing was performed in MeshLab (v2020) [17], and the final 3D rendering was carried out in ParaView (version 5.8) [18].

For BMP9-KO samples, the following process was used. In the Pixel Classification workflow, the segmentation is generalized by Batch Processing. This allows the trained classifier to be applied to the dataset. Manual corrections were subsequently performed using Microscopy Image Browser (MIB). The 3D mesh is then generated in Avizo. Finally, the 3D mesh can be visualized and explored using ParaView.

##### nnU-Net

The Python package nnU-Net [12] is a popular self-configuring tool for deep learning-based biomedical image segmentation. It automatically adjusts preprocessing, network architecture, training, and post-processing parameters, according to the input data. This makes it particularly interesting for a semi-automatic segmentation workflow.

Starting from the generated ground truth, nnU-net was used to obtain clean segmented volumes and generalize segmentation to larger or previously unseen raw volumes. Ilastik’s segmentations were used as “Ground Truth” to train nnU-Net’s underlying CNN model.

nnU-Net uses five-fold cross-validation for hyperparameter tuning, dividing training data into five subsets and repeating training five times (**Figure S5**). With nnU-Net, users can still choose a few different parameters, including the type of model (e.g., standard U-Net vs residual encoder U-Nets, etc), and its dimensionality (i.e., 2D vs 3D). When working with large volumes, it is also advisable to divide them into smaller sub-volumes. This allows nnU-Net to organize the sub-volumes into batches, resulting in better use of the GPU memory and faster convergence of the optimizer. Moreover, smaller volumes require smaller receptive fields for the segmentation model, thus reducing the model’s size and the inherent risks of overfitting. When taking the sub-volume slicing into account, for our workflow, the key parameters include: dimensionality (2D vs 3D U-Net selection based on computational resources and data complexity), sub-volume size (balancing spatial context preservation with sufficient sub-volume representation), using overlap to prevent boundary discontinuities, **epochs** (150 for 2D, 350 for 3D U-Net convergence, **Figure S6**), and optimizer (Adam and SGD algorithms). The most effective approach for improving segmentation predictions on a new dataset is fine-tuning. The user manually annotates a small set of consecutive images, typically around ten, selected from a randomly chosen image stack. A model previously trained on a ground truth is then further trained to adapt to the new data through rapid segmentation refinement. Once fine-tuned, the model generalizes the segmentation predictions to the entire 2D image stack of the new dataset.

##### Quantitative analysis of LSEC fenestrations

###### Fenestrae counting

The overall methodology is described in **Figure 6**. The 3D binary mask of LSEC was converted to a 3D mesh using Avizo (Thermo Scientific). The original mesh containing millions of faces was then simplified in MeshLab [17] to reduce the number of faces to about 200,000 faces. Coordinates of the mesh vertices were scaled into [0, 1] and ISOMAP [19] was applied to embed the mesh into 2D. To reduce the computation, 30,000 vertices among all vertices were randomly selected and used in ISOMAP to learn the mapping function. We next rescaled the embedded 2D mesh to the original scale and rasterized it into a 2D binary image. We used morphological opening with a disk size of 3 pixels to correct the rasterization artifact. Distinct objects were then identified in the image, and the pore-objects were selected based on an object area from 50 pixels^2^ to 10,000 pixels^2^. The pores touching the image border were not counted. The pipeline requires an open 3D mesh. In case of a closed mesh, we used MeshLab to manually identify a fenestration-free region as the nuclear region and to truncate the mesh at this location, thereby generating an open mesh. The LSEC fenestration counting was implemented in the Python library GeNePy3D [20] and the source code is available at https://lsec-fenestrae-counting-8b6ca1.pages.pasteur.fr/README.html.

###### Fenestrae diameter distribution

The distribution of LSEC fenestration diameters was analyzed using the cPSD (continuous Pore Size Distribution) method from the Fiji plugin *xlib*, developed by Beat Münch [21]. The cPSD technique provides an accurate quantification of pore sizes in the form of a cumulative distribution [22]. The pore space is modelled as a continuous volume filled with spheres of varying radii [21]. The cPSD method allowed the measurement of fenestration radii throughout the entire volume, over a radius range of 20-150 nm. This interval was selected based on literature reports describing LSEC fenestration diameters between 50 and 300 nm [23]. A step size of 4 was used, corresponding to the pixel size in our case. The analyses were performed on 3D reconstructions of sinusoids. Once the cPSD simulation was complete in Fiji, a text file was generated containing three columns: fenestration radius (nm), number of voxels greater than or equal to the radius, and pore volume (nm³). Calculations in Excel, along with normalization, were then applied to determine the frequency distribution of fenestrations radii.

## Results

### Sample preparation with optimal preservation and differential contrast for LSEC

To preserve liver architecture as best as possible, and in particular sinusoid ultrastructure and LSEC fenestrations [9], liver tissue was initially fixed by mice perfusion followed by high-pressure freezing of small liver pieces (∼1 mm^3^). The main improvement over the en-bloc staining method and conventional methods was that staining and water substitution were performed in cryogenic conditions. The optimized protocol resulted in LSEC membranes appearing darker than the underlying hepatocyte microvilli (**Figure 1A-B**), which was not the case with the conventional preparation protocol (**Figure 1C-D**). This improvement enables selective segmentation of LSEC, which is a prerequisite for the development of a methodology enabling to analyze accurately LSEC fenestrations based on FIB-SEM data.

**Figure 1.**
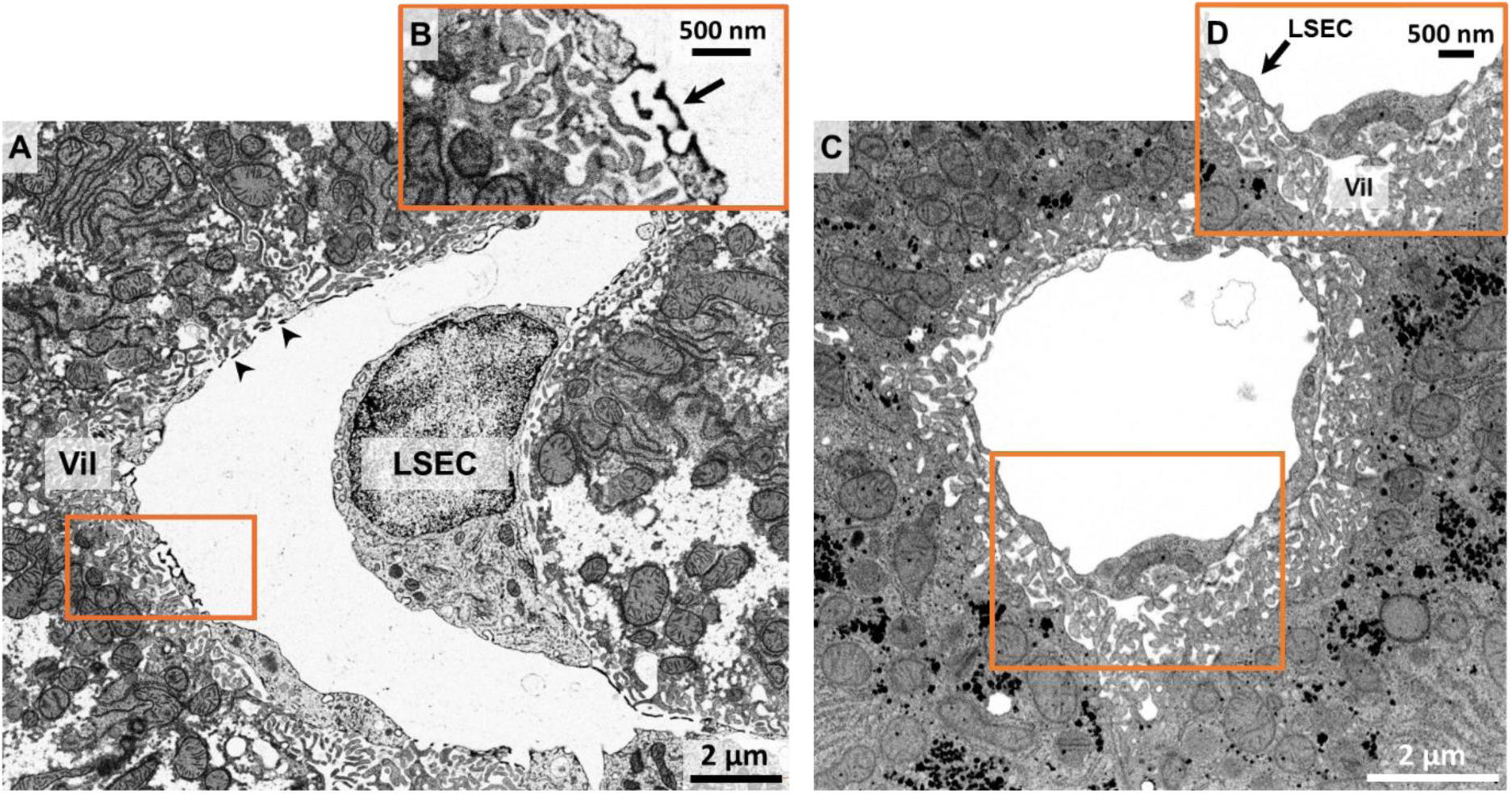
Comparison of sample staining from FIB-SEM data. A, Single FIB-SEM image from a liver block prepared with the optimized sample preparation protocol, with zoom-in, in **B**, highlighting the differential contrast between LSEC (darker, black arrow) compared to the underlying hepatocyte microvilli. **C**, Single FIB-SEM image from a liver block prepared without high-pressure freezing nor cryo-substitution, with zoom-in, in **D**, on a LSEC membrane that does not exhibit a higher contrast than underlying hepatocyte microvilli.

### Strategy for LSEC ground truth segmentation

FIB-SEM acquisitions of hepatic sinusoids were performed with an isotropic resolution of 4 × 4 × 4 nm³. However, at this resolution, the generated image stacks are very large, often reaching several hundreds of gigabytes. This makes them difficult to process and segment. Binning the data reduced the file size, but led to the loss of some fenestrae during segmentation (**Figure S7**), indicating that correct segmentation of LSEC fenestrations with minimal artifacts requires working with the initial data at 4 × 4 × 4 nm³ resolution. The membrane-based carving segmentation method [24]in ilastik failed to segment LSEC due to the large number of annotations required to delineate fenestrations, where the LSEC membranes become very thin, and because of the difficulty in distinguishing these thin structures from the surrounding tissue in the space of Disse, including hepatocyte microvilli and stellate cell cytoplasmic extensions. These neighboring structures exhibit similar shapes with LSEC, creating substantial segmentation noise. Additionally, LSEC occupy a narrow zone relative to the surrounding hepatocytes, which further complicates the segmentation task. This one-step segmentation approach proved unsuitable for LSEC segmentation, as it exceeded the workstation memory and ultimately failed.

Taking into account the LSEC shape which comprises a large nuclear region and thin fenestrated regions, we developed an optimized multi-step segmentation workflow for LSEC segmentation in 3D (**Figure 2**). To address challenges related to LSEC shape and position, we designed a four-step cascade segmentation approach that separates the sinusoid edge from the neighboring cellular structures while handling each LSEC’s region distinctly. The first step defines the LSEC “fenestration localization region”, which highlights the thin zone between the sinusoidal lumen and the space of Disse. This region was extracted by performing a coarse sinusoidal lumen segmentation on heavily binned data, followed by outline extraction, dilation, and Gaussian filtering The LSEC fenestrated regions were segmented in the second step using the pixel classification method (in ilastik) [25] with a dual-channel input combining the original images and the fenestration localization region. The latter serves as a spatial prior, guiding the classifier to focus on the narrow zone containing fenestrations.

**Figure 2.**
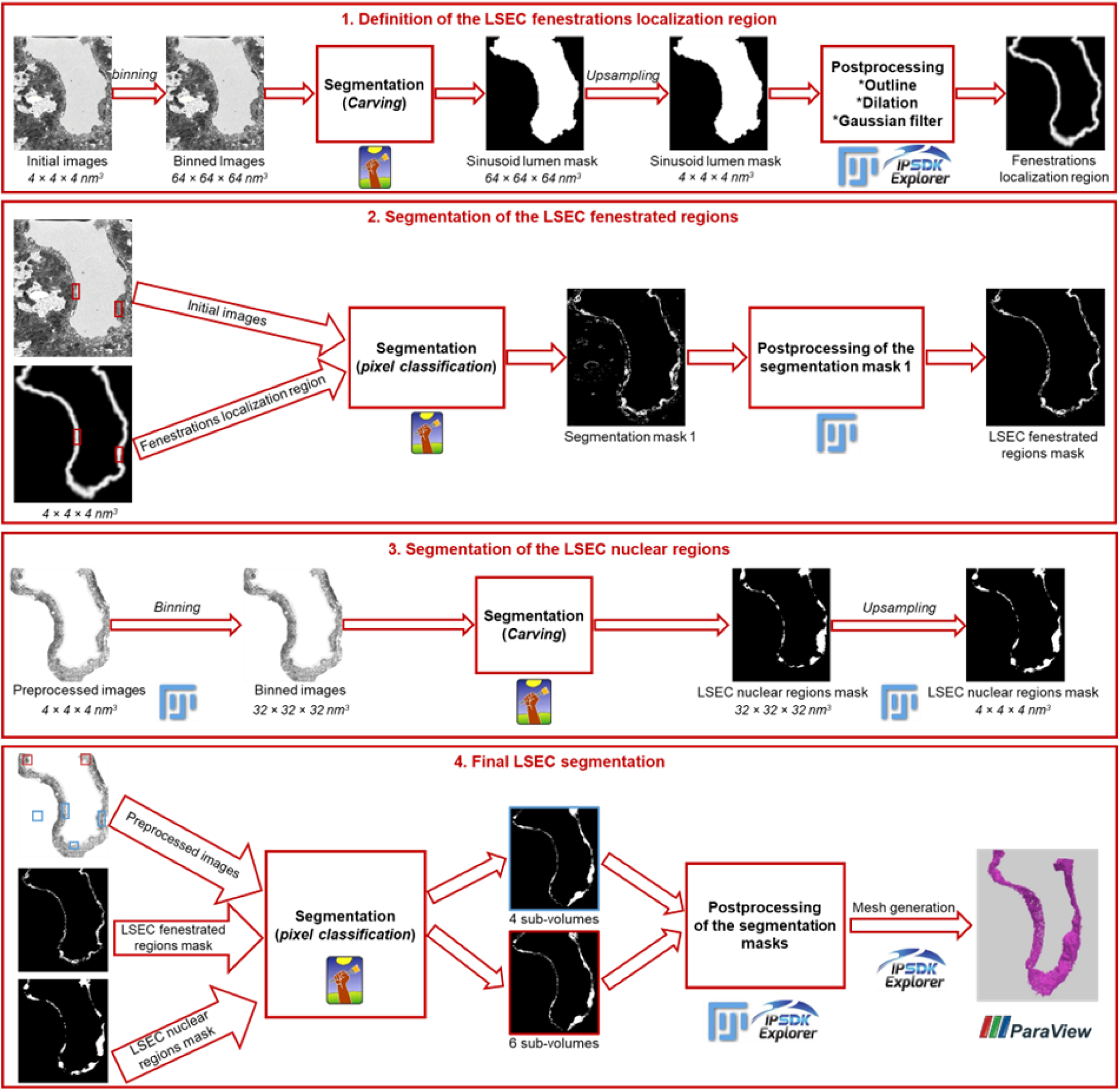
Multi-step LSEC segmentation workflow. In the final step, the 4 sub-volumes used in the training are shown in blue, with the 2 additional sub-volumes shown in red, bringing the total to 6.

Training on selected sub-volumes enables segmentation generalization to the entire image stack. Post-processing using a subtraction mask corrected the residual segmentation errors in hepatocytes.

In the third step, LSEC nuclear regions were segmented using the carving method. Since hepatocyte membranes introduce noise in the boundary map, the images were pre-processed by defining an LSEC localization region that encompasses both fenestrated and nuclear regions while minimizing hepatocyte microvilli inclusion.

This involves determining LSEC extensions towards the sinusoidal lumen (LSEC luminal region) and towards the hepatocytes (LSEC mesenchymal region) through controlled dilation and subtraction operations, generating a mask that reduces hepatocyte interference during segmentation.

Finally, segmentation results from steps two and three were combined using a pixel classification segmentation with a three-channel input: the preprocessed initial images, the LSEC fenestrated region mask, and the LSEC nuclear region mask. Training on different sub-volume combinations (4 and 6 sub-volumes, shown in blue and red in **Figure 2**) optimizes segmentation of different LSEC regions, with nuclear regions better segmented using 4 sub-volumes and fenestrated regions using 6 sub-volumes. The segmentation masks resulting from both trainings are combined to generate the final mask used to reconstruct the 3D mesh of the sinusoid.

This cascade workflow successfully achieves 3D LSEC segmentation while preserving fenestrations. It leverages the complementary strengths of ilastik’s carving and pixel classification methods while circumventing their individual limitations. The carving method efficiently segments membrane-delimited structures, whereas the pixel classification method enables to train the classifier on small sub-volumes and to generalize the results to large datasets through its batch processing option. Furthermore, strategic use of binning and resampling addresses computational constraints without compromising the segmentation quality. The resulting 3D mesh clearly visualizes both broad LSEC nuclear regions and thin fenestrated regions (**Figure 3** and **Video S1**), enabling detailed ultrastructural analysis of hepatic sinusoids.

**Figure 3.**
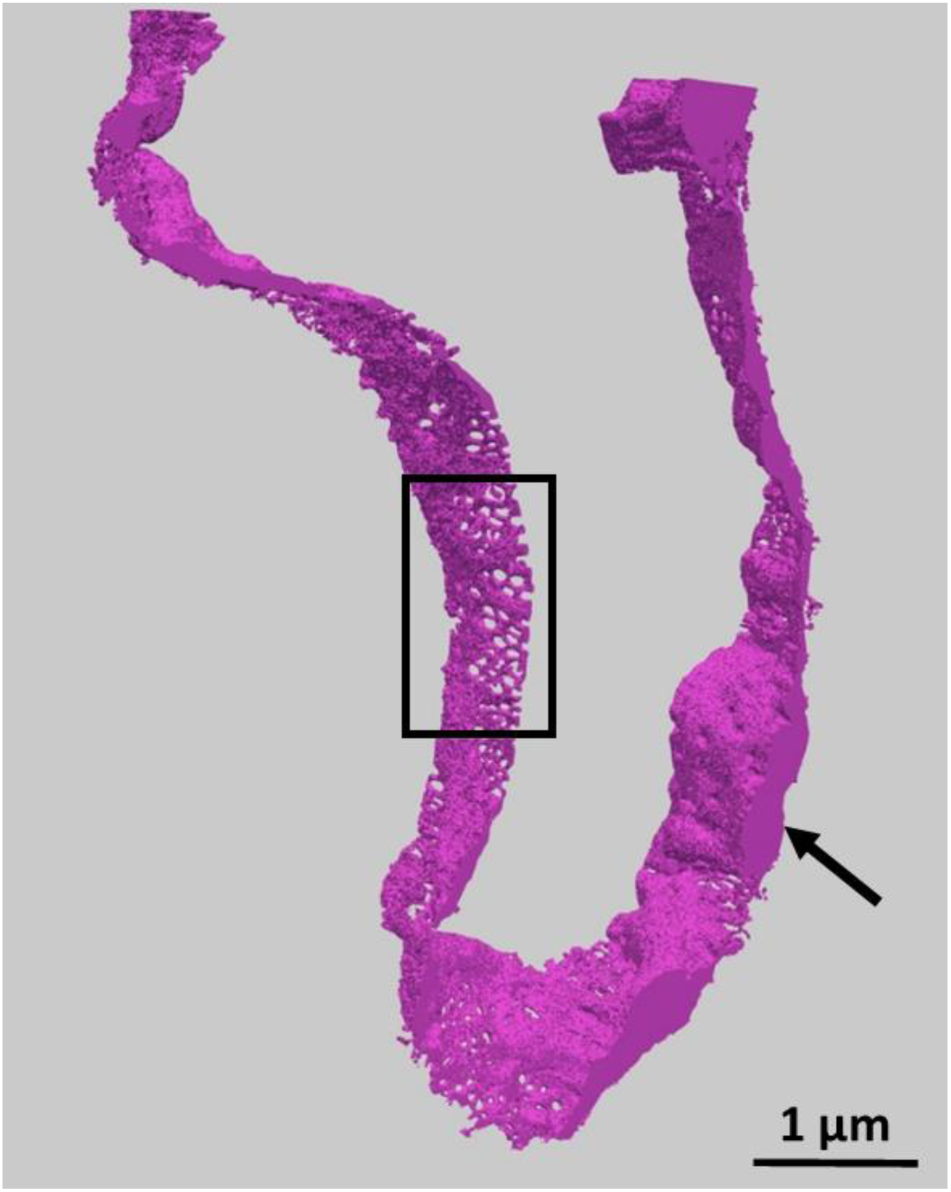
Final LSEC segmentation obtained using the multi-step workflow. The black arrow highlights a wide nuclear region and the black box a thin fenestrae region.

### Towards semi-automatic LSEC segmentation

The optimal LSEC segmentation obtained was used as a ground truth for deep learning training in nnU-Net. This is necessary for scaling the segmentation to larger and multiple datasets. This, in turn, enables obtaining enough data to perform statistical analysis on fenestrae numbers and diameters. The main advantage of nn-UNet is that it can automatically determine suitable configurations for pre- and post-processing and network architecture. Another key benefit of this tool is that users can directly interact with and control the model behavior by adjusting specific hyperparameters, leading to more robust segmentation results. This is particularly important when dealing with complex objects containing both large regions and thin structures. To obtain reliable statistical data, it is also important not to limit the batch-processing step when propagating segmentation across FIB-SEM slices, as is the case in ilastik.

For the correct execution of this step, the following hyperparameters were optimized: the patch size was set to (30, 900, 900) for the WT sample, corresponding to the number of pixels along (Z, X, Y) with an overlap of (50, 50, 50) pixels between patches. The chosen optimizer was Adam, with a learning rate of 0.001 and cosine annealing scheduling. In the 2D U-Net configuration, the number of epochs was set to 150, whereas in the 3D U-Net configuration, it was set to 350. The higher number of epochs in 3D is required, due to the increased number of parameters and the inherent complexity of volumetric features. After an optimal training, the segmentation of the same dataset using nnU-Net led to several improvements over the one performed in ilastik. Segmentation errors were removed, the LSEC outlines are better defined and the objects become continuous thanks to the removal of isolated pixels wrongly annotated either in the object or in the background (**Figure 4**).

**Figure 4.**
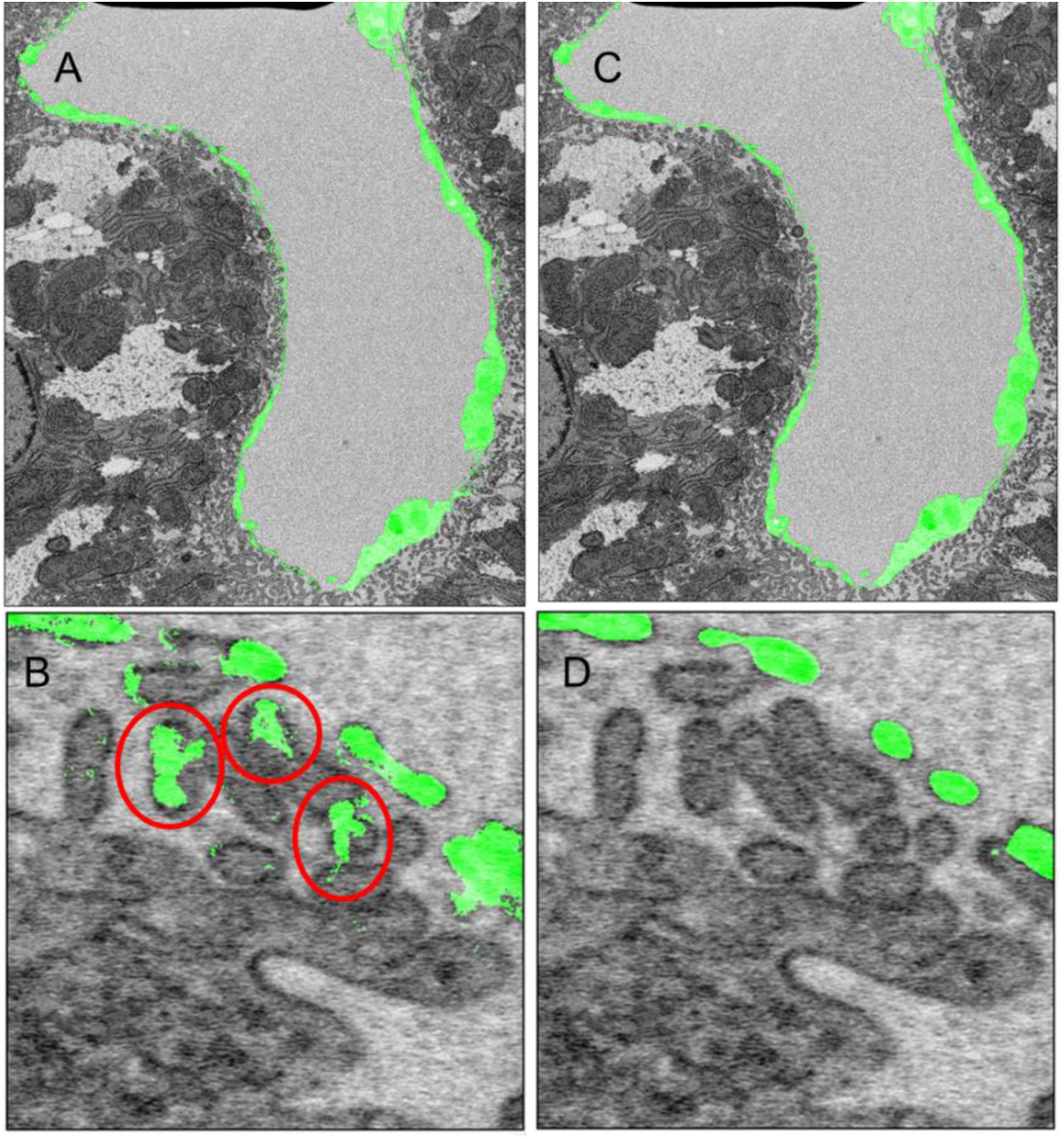
nnU-Net segmentation improvement. Segmentation performed with ilastik, large field of view (**A**) and zoom-in (**B**) showing improperly assigned hepatocyte microvilli (red circles). Segmentation of the same region obtained with nnU-Net (**C-D**).

### Quantification of LSEC fenestration

Current methods for determining LSEC fenestrae densities and diameters are mainly manual, precluding the possibility to perform large scale comparative and statistical analyses. To be consistent with the development of a pipeline for LSEC fenestration analysis, a semi-automatic quantitative method is required. To this end, we implemented a dual approach using, on the one hand, cPSD to determine diameter distribution in the analyzed LSEC segment, and on the other hand, we developed a tool based on manifold embedding allowing us to count the number of fenestrae in the same segment. The cPSD method enables the measurement of pore sizes throughout the 3D volume, using a user-defined range and a step size. Originally developed for analyzing porous materials, its application to biological tissues is novel. One of the key advantages of cPSD is that it is fully automated, ensuring statistically robust and unbiased measurements, independent of manual intervention. For our analysis, used the 3D variant of the cPSD method, as our data consisted of a stack of images.

The distribution of fenestrae diameters is gaussian (**Figure 5**) with average and median diameters of 142 and 140 nm, respectively. These data are consistent with the literature [6] confirming the suitability of this method.

**Figure 5:**
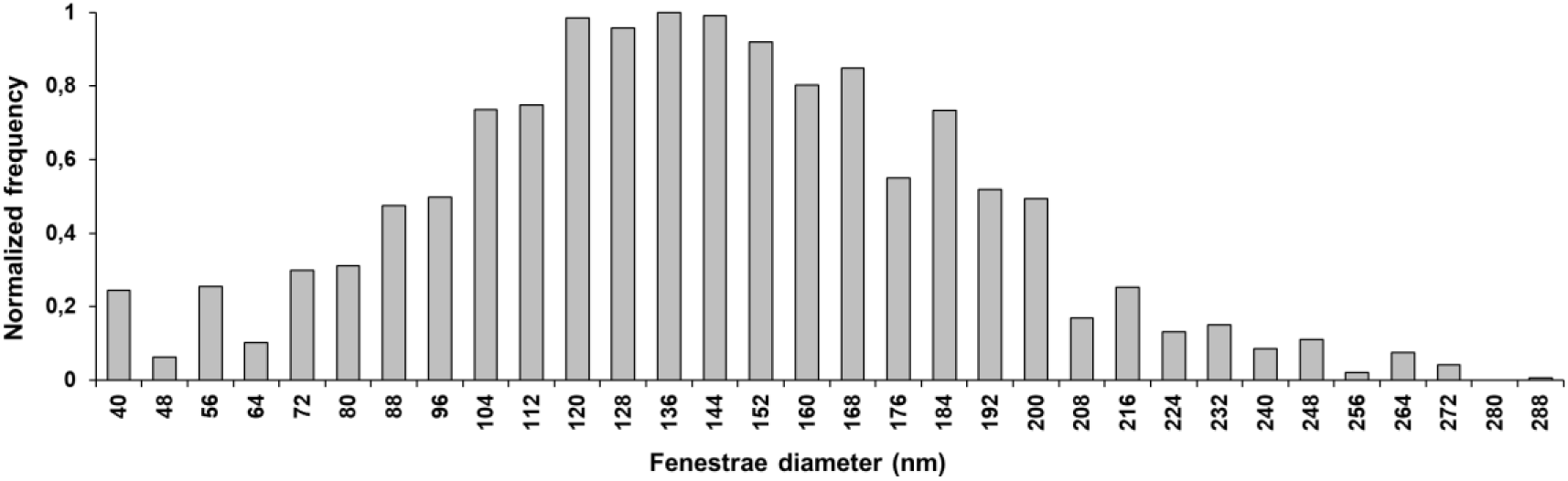
Distribution of LSEC fenestrae diameters in the WT dataset. Automated measurement method for determining LSEC fenestrae diameters from the binary segmentation mask, using the continuous Pore Size Distribution (cPSD method) under the xlib plugin in Fiji, normalized histogram. Measurements were performed on data with a voxel size of 4 × 4 × 4 nm³.

Since the cPSD approach does not provide individual diameters, it cannot provide counts of fenestrae. Moreover, automatic pore counting in 3D is not possible. Therefore, to determine the number of fenestrae per LSEC segment, we developed a method using ISOMAP [19] to embed the 3D mesh of LSEC into 2D. Then, the 2D embedded mesh was converted to an image and the pores, that corresponds to fenestrae, were identified by labelling the distinct objects in the image (**Figure 6**). On the WT dataset, a total of 168 fenestrae were counted, corresponding to a density of 10.8 fenestrae per µm². These data are in good accordance with previously published data showing around 6 fenestrae per µm² [6].

**Figure 6:**
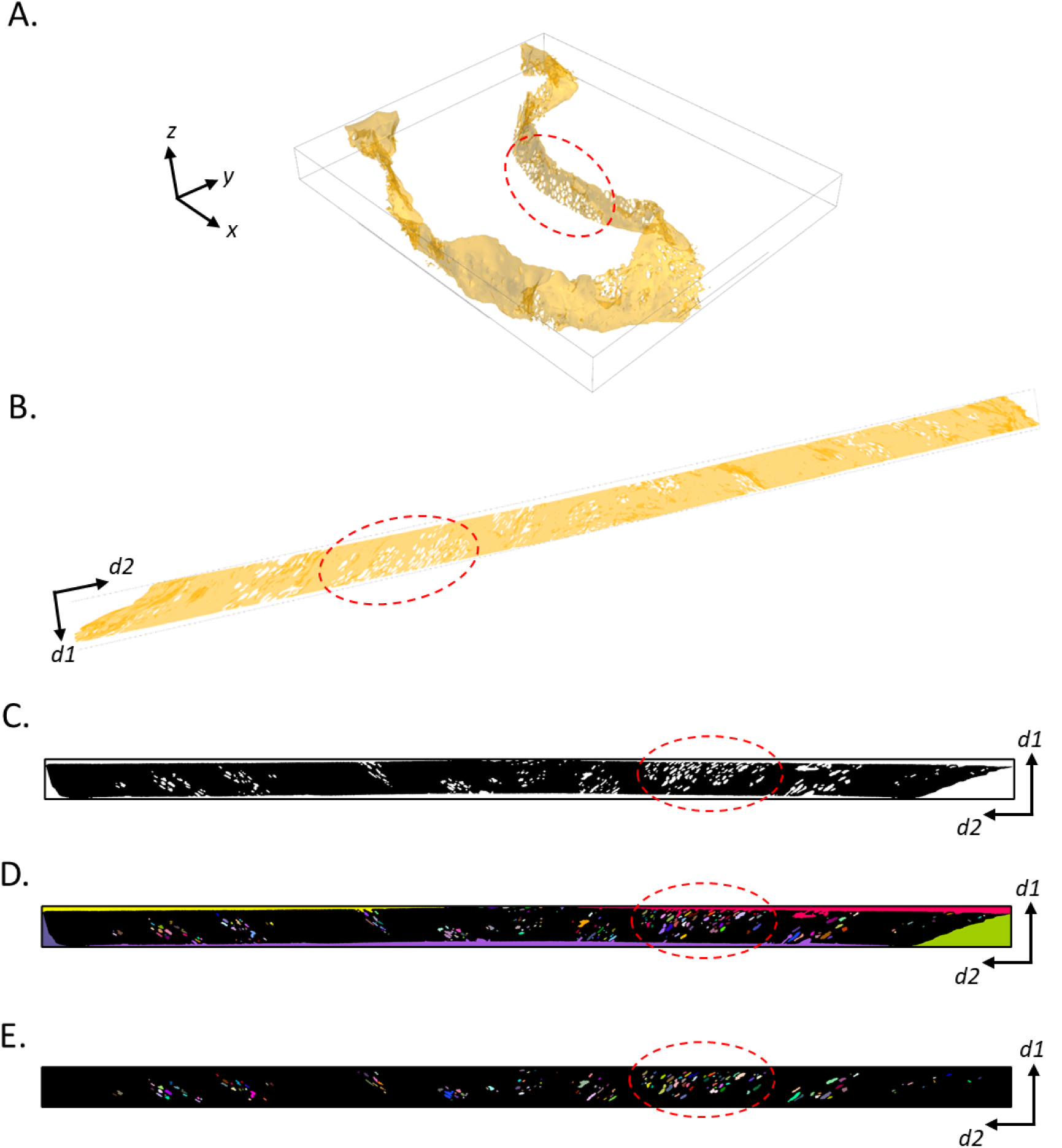
LSEC fenestrae counting. **A**, 3D mesh of LSEC generated from its 3D binary mask. It is simplified and smoothed in MeshLab. **B**, 2D embedding of 3D mesh using ISOMAP. The 2D mapping preserves the geodesic distances on the 3D mesh. **C**, Image rasterization from 2D embedded mesh. Morphological opening is used to correct the rasterization artifact. **D**, Image labelling to identify distinct objects in the image. **E**, Removal of undesired objects based on the object area. The pores touching the image borders are not counted. The dashed circle in red highlights a LSEC region throughout the steps as a reference.

### Generalization of the pipeline to liver with altered LSEC ultrastructure

Since our pipeline proved to be efficient for LSEC segmentation and quantitative analysis of their fenestration, we decided to apply it to other types of liver samples with known alterations in LSEC ultrastructure. It has been shown that BMP9 is important for LSEC differentiation and fenestration maintenance, and its deletion leads to alteration in LSEC fenestration [6]. Liver samples from mice with genetic ablation of *Bmp9* were thus imaged by FIB-SEM. The 3D reconstruction of the sinusoid appeared different compared to samples from WT mice. There is a lower density of fenestrae with more variable diameters, including large ones (**Figure 7A**, white arrow). Initially, segmentation was performed with nnU-Net using the training from WT mice samples. However, this led to a limited segmentation of the LSEC missing some fenestrated parts (**Figure 7B**, red circle). This is likely due to the fact that the object we want to segment in the *Bmp9-KO* dataset has features different from those in the WT sample. To obtain a better segmentation of the *Bmp9-KO* images, a specific segmentation ground truth was built (**Figure 7C**). FIB-SEM data shows higher contrast for LSEC membranes in WT sample compared to *Bmp9*-KO, enabling segmentation with pixel classification in ilastik using only two classes, the object (LSEC) and the background. In contrast, *Bmp9-KO* required three classes to better guide the network during training, as the grayscale contrast was less pronounced, and the LSEC membranes were more discontinuous. As for the WT, the ground truth generated in ilastik was improved after nnU-Net has been trained on it leading to more complete and accurate segmentation of the LSEC, in particular, of the fenestrated region (**Figure 7D**, red circle).

**Figure 7:**
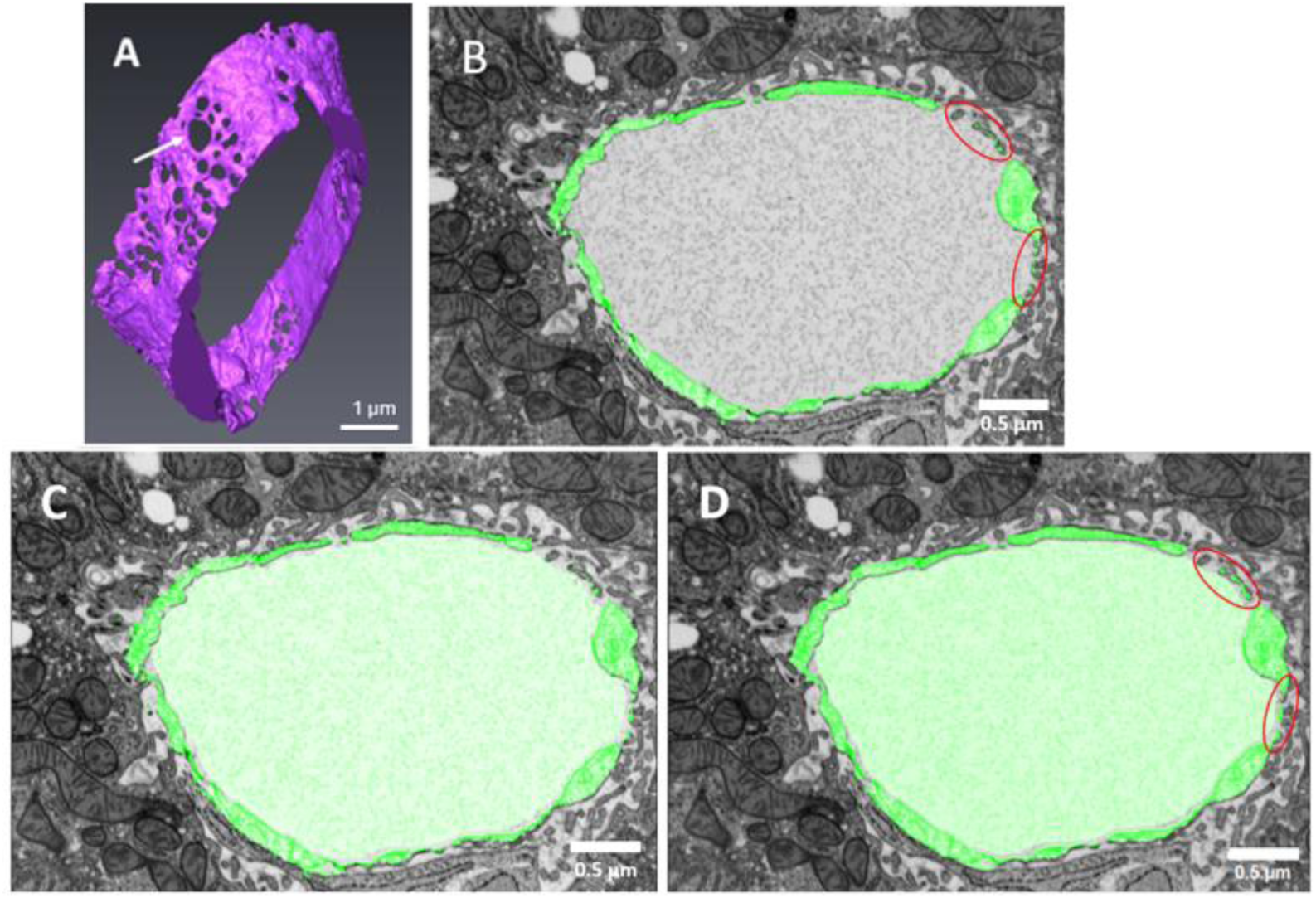
Segmentation of a sinudoid from a Bmp9-KO mouse liver. (**A**) 3D reconstruction of a Bmp9-KO hepatic sinusoid showing large fenestrae (white arrow). (**B-C**) Ilastik-based segmentation of LSEC membranes using a method similar to that used for WT sample with two classes (**B**) or using three classes (**C**). (**D**) Prediction results obtained with nnU-Net based on the three-class ground truth.

Using LSEC segmented volume from the *Bmp9-KO* dataset, diameter distribution and fenestrae counting were determined using cPSD and manifold embedding methods, respectively. Contrary to WT samples, fenestrae diameter distribution appeared to be non-Gaussian (**Figure 8**), making average diameter not relevant. The median diameter of the distribution is 160 nm. In the KO dataset, a total of 53 fenestrae has been counted, leading to an average of 6.5 fenestrae per µm². Therefore, the diameter is slightly larger and the density of fenestrae is lower in *Bmp9-KO* compared to WT LSEC. These results are in agreement with data previously published on the same mice [6]. Overall, using an appropriate ground truth, LSEC can be accurately segmented from various samples, allowing reliable quantification of fenestrae diameter distribution and density.

**Figure 8:**
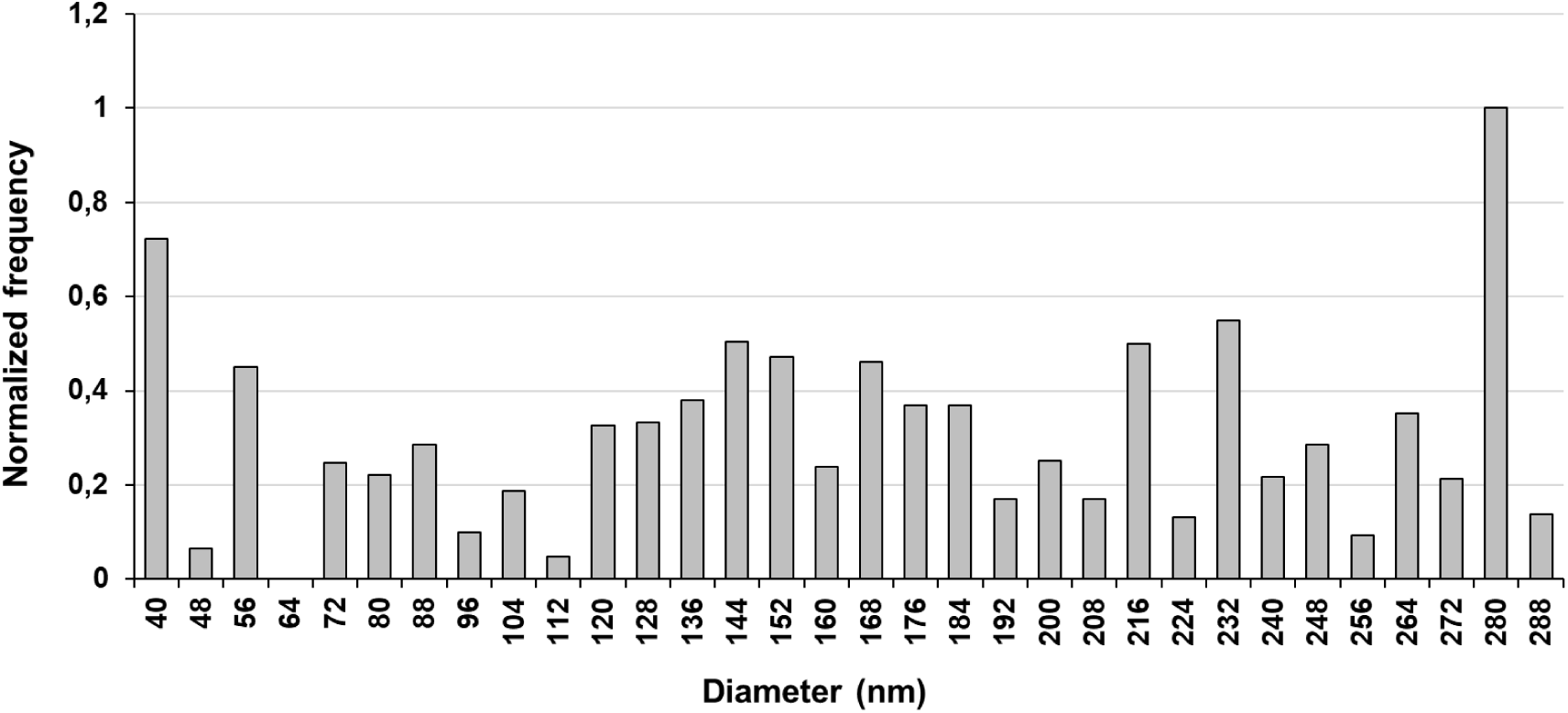
Distribution of LSEC fenestrae diameters in BMP9-KO dataset. Automated measurement method for determining LSEC fenestrae diameters from the binary segmentation mask, using the continuous Pore Size Distribution (cPSD) under the xlib plugin in Fiji, normalized histogram. Measurements were performed on data with a voxel size of 4 × 4 × 4 nm³.

## Discussion

In this study, we developed a complete workflow enabling to reconstruct the fenestrated endothelium of LSEC from mouse liver in 3D and at nanometer scale, and to quantify the number and the diameter distribution of the fenestrae in two mouse phenotypes. Given the critical role of the fenestrated endothelium in liver functions, this methodology is of paramount interest in the field of liver biology. We have shown that the methodology can be applied to samples with altered LSEC ultrastructure. It can therefore be used in various contexts to characterize the fenestrated endotheliumupon aging or in liver-related diseases. Indeed, herein, our methodology was used to compare WT and *Bmp9-KO* mice livers. We showed that the distribution of fenestrae diameter is more heterogenous in *Bmp9*-*KO* compared to WT sinusoid endothelium, with a diameter distribution that clearly shows a significant number of fenestrae with diameters larger than 200 nm in *Bmp9-KO* data, which are almost absent in the WT dataset. Moreover, the density of fenestrae in the analyzed *Bmp9-KO* data is higher, and the fenestrated endothelium visually appears less affected in our FIB-SEM data compared to previously published SEM data [6]. This difference could be due to the type of sample analysed: liver tissue in a near-native state in the current study, compared to LSEC isolated from the liver and cultured in petri dishes in the previous study [6].

Mouse liver was used as a test case, but the methodology could be extended to other species, particularly humans, in order to identify liver dysfunction or monitor disease progression. Indeed, including nnU-Net in the workflow makes it very powerful and generalizable to various samples, with the ability to process large volumes. The originality of our study lies in the application of nnU-Net to a specific cell type within the liver and not to an entire organ, as is more commonly done [26]. Although ground truth segmentation is assumed error-free, sparse annotations often introduce inaccuracies. nnU-Net predictions demonstrate the capacity to correct these, highlighting its robustness. In our study, the fine-tuning strategy yielded the most accurate predictions. This approach relies on adapting a pre-trained model rather than training from scratch. In terms of the time a user would have spent segmenting the dataset using only ilastik, it would have taken approximately one week of annotations due to the complexity of the LSEC which include both thin structures and larger regions. nnU-Net thus leads to important time saving. In addition, nnU-Net offers high precision, as it allows the adjustment of hyperparameters specific to our dataset, helping to prevent overfitting and providing the most possible accurate segmentation. In conclusion, this powerful workflow developed for the analysis of the liver fenestrated endothelium can be applied to a wide range of liver samples, including diseased tissues, especially from human patients.

## Acknowledgments

The authors are grateful to Soumalamaya Bama and Charlène Magallon for their help in the maintenance of mouse colony, Patrice Perrenot and Thomas David for their help in segmentation tools implementation, and Kassem Dia and Matthew Bryan for their help with nnU-Net set up.

This work used the platforms of the Grenoble Instruct-ERIC Center (ISBG: UMS 3518 CNRS-CEA-UGA-EMBL) with support from FRISBI (ANR-10-INSB-05-02) and GRAL (ANR-10-LABX-49-01) within the Grenoble Partnership for Structural Biology (PSB). The EM facility is headed by Guy Schoehn and supported by the Auvergne Rhône-Alpes Region, the Fondation Recherche Medicale (FRM), the fonds FEDER and the GIS-Infrastrutures en Biologie Santé et Agronomie (IBISA). Part of this work was carried out on the Platform for Nanocharacterisation (PFNC) of CEA Grenoble, supported by the “Recherche Technologique de Base” and “France 2030 - ANR-22-PEEL-0014” programs of the French National Research Agency (ANR). We acknowledge France-BioImaging infrastructure (https://ror.org/01y7vt929) supported by the French National Research Agency (ANR-24-INBS-0005 FBI BIOGEN).

## Notes

*Financial support and sponsorship.* YR and CP PhD as well as the animal facility platform were supported by the LabEx Gral, the Grenoble Alliance for Integrated Structural & Cell Biology, a program from the Chemistry Biology Health Graduate School of University Grenoble Alpes (ANR-17-EURE-0003). This work was also supported by the UGA IRGA funding (project 3DEM-Liver), the LabEx Arcane (ANR-17-EURE-0003), the Ambition Internationale program from the Région Auvergne Rhône-Alpes, the Institut National de la Santé et de la Recherche Médicale (INSERM), the fondation pour la recherche médicale (EQU202003010188) and the National Infrastructure France-BioImaging (https://ror.org/01y7vt929) supported by the French National Research Agency (ANR-24-INBS-0005 FBI BIOGEN).

Conflicts of interest: nothing to report

### Competing Interest Statement

The authors have declared no competing interest.

