## Supplementary methods and figures for "A general methodology for liver sinusoid fenestration analysis based on 3D electron microscopy data"

### LSEC segmentation full method

#### *Step 1: Definition of the LSEC fenestration Localization Band*

The initial image stack is first binned in Fiji (using the *Average* method with a *binning* factor of 16 in each spatial direction). The resulting image stack has the following dimensions: $231 \times 268 \times 25$ px³ with a voxel size of $64 \times 64 \times 64$ nm³. This binned image stack is then used as *Raw Data* for an Ilastik *(Carving)* project for sinusoid lumen segmentation. The following parameters are used: *dark lines*, filter scale = 1.6. The segmentation mask exported from Ilastik is resampled in Fiji using bicubic interpolation with a scaling factor of 8 in all 3 directions.

The post-processing steps for this resampled mask are then performed in IPSDK, which is an image analysis library written in Python and commercialized by the company Reactiv'IP (https://www.reactivip.com/fr/traitement-dimages/). These steps are performed in the following order:

- 'Binarization' of the mask (necessary step even though the segmentation mask is initially binary) (*Binarization* ⟶ *Simple Threshold*)
- Application of the *outline* filter (*Basic Morphology* ⟶ *Boundary 3D*) to extract the sinusoid contour. The 6-nearest neighbor connectivity of a voxel (*connectivity 6*) is selected in *Neighborhood*
- Application of 3D dilation (*Basic Morphology* ⟶ *Dilation 3D*) with a radius of 30 using a spherical structuring element

Finally, a Gaussian filter with variance (*sigma*) of 30 in all 3 directions is applied in Fiji: *Process* ⟶ *Filters* ⟶ *Gaussian Blur 3D*. Thus, the fenestration mask is created.

#### *Step 2: Segmentation of LSEC Fenestrated Regions*

The initial image stack and the fenestration mask (from step 1) are combined in Fiji to form a 2-channel image stack. Two sub-volumes of dimensions 354 x 393 x 392 px³ are extracted from this image stack and converted to HDF5 format in Fiji before being imported as Raw Data into an ilastik project (pixel classification). All features proposed by ilastik are selected. After segmentation of these two sub-volumes, the segmentation is generalized to the entire (2-channel) image stack: in the batch processing tab of the ilastik project, click on Select Raw Data Files, import the entire image stack to be segmented (2-channel image stack) and finally click on Process all files. To export the binary segmentation mask and not the probability map, choose Simple Segmentation in the Prediction Export tab (as explained above).

After batch processing, the generated binary mask (segmentation mask 1) is post-processed in Fiji by applying the following steps:

- Inversion of the fenestration mask followed by 3D dilation with radius 50 using a spherical structuring element. Since IPSDK is faster than Fiji for image analysis, it is used for dilation. The radius is chosen after trial-and-error steps. The resulting mask is called the subtraction mask.
- Subtraction of the subtraction mask from the mask generated after batch processing: Process → Image Calculator, put segmentation mask 1 in Image1, the subtraction mask in Image2 and Subtract in Operation. Both image stacks must be open in Fiji before performing the subtraction.

The final result of this step is the LSEC fenestration segmentation mask.

***Step 3: Segmentation of LSEC Nuclear Regions***

***Initial Data Preprocessing***

**Determination of the sinusoid outer edge**. In IPSDK, the resampled segmentation mask of the sinusoid lumen (see step 1) is dilated in 3D with radius 285 using a spherical structuring element. Then, the non-dilated mask is subtracted from the thus dilated mask (Arithmetic → Subtraction Image). The result defines the sinusoid outer edge.

**Determination of the sinusoid inner edge.** The following steps are performed in IPSDK:

- Dilation of the resampled segmentation mask of the sinusoid lumen with radius 85 using a spherical structuring element
- Subtraction of the non-dilated mask from the dilated mask. The result is called the intermediate mask.
- Dilation of the sinusoid contour (see step 1) with radius 85 using a spherical structuring element.
- Subtraction of the intermediate mask from the dilated contour mask. The result defines the sinusoid inner edge.

**Definition of the LSEC localization band.** The two inner and outer edges are added in Fiji: Process → Image Calculator, choose Add in Operation and put the two image stacks representing the two sinusoid edges in Image1 and Image2. Both image stacks must be open in Fiji before performing the addition.

**Application of the LSEC localization band to initial images.** The following steps are performed in Fiji:

- Subtraction: LSEC localization band - initial data
- Contrast inversion of the subtraction result

***Segmentation of LSEC Nuclear Regions***

The preprocessed initial images are binned in Fiji using the Average method with a binning factor of 8 in all 3 directions. The resulting image stack has the following dimensions: $462 \times 535 \times 49$ px³ and a voxel size of $32 \times 32 \times 32$ nm³. It is then used as Raw Data for an Ilastik project (carving) with parameters dark lines, filter scale = 1.6. The binary segmentation mask is exported from Ilastik and resampled by interpolation in Fiji. This mask is the LSEC nuclear region mask.

***Step 4: Final LSEC Segmentation***

The preprocessed initial images (voxel size = 4 × 4 × 4 $nm^3$), the LSEC fenestration mask, and the LSEC nuclear region mask are combined in Fiji to form a 3-channel image stack: Image → Color → Merge Channels. In total, 6 sub-volumes are extracted from this image stack and converted to HDF5 format in Fiji before being imported as Raw Data into an Ilastik project (pixel classification).

All features proposed by Ilastik are selected. After training the classifier on the sub-volumes, the segmentation is generalized to the entire (3-channel) image stack. The dimensions of the sub-volumes used in algorithm training are detailed in **Table S1**.

| **Sub-volume number** | **1** | **2** | **3** | **4** | **5** | **6** |
|---|---|---|---|---|---|---|
| **Dimensions ($px^3$)** | 160×242 ×392 | 171×474 ×392 | 216×216 ×392 | 231×188 ×392 | 262×292 ×392 | 286×321 ×392 |

**Table S1: Dimensions of the sub-volumes used for Pixel classification algorithm training in step 4 of hepatic sinusoid segmentation.**

The binary masks of the entire image stack obtained after training on 4 sub-volumes (sub-volumes 1 to 4 from **Table S1**) and on 6 sub-volumes (sub-volumes 1 to 6 from **Table S1**) are exported from Ilastik. Then, after conversion to TIFF format, they are post-processed in Fiji by applying the OR arithmetic operation: *Process → Image Calculator*, put the two masks as *Image1 and Image2* and choose *OR* in *Operation*. As mentioned above, both image stacks must be open in Fiji before performing this operation. The resulting mask is then imported into IPSDK. To reduce segmentation errors of this binary mask, a connected components analysis is initially applied to separate its connected components (groups of connected voxels):

*Advanced Morphology* → *Connected Component 3D* and choosing 6-nearest neighbor connectivity of a voxel (*connectivity 6* in *Neighborhood*).

Size filtering is then performed on the detected components (or objects) to keep only the largest component: *Advanced Morphology* → *Keep big shape 3D*. The corresponding 3D mesh is created: *Shape Segmentation* → *Label Shape Extraction 3D*, in *Label Image*, put the result obtained after the *Keep big shape 3D* operation. The generated 3D mesh is saved in the *Values* → *Shapes* menu (top left). It is saved in STL format (*Standard Triangle Language*, with .stl extension) and converted to OBJ format in Paraview. Then, it is simplified in MeshLab: Filters → *Remeshing, Simplification and Reconstruction* → *Quality Edge Collapse Decimation*. The chosen *Quality threshold* parameter is 0.3 to reduce the number of mesh surfaces by 30%, the *Optimal position of simplified vertices* option is checked, and other parameters are left at their default values. The simplified mesh is finally smoothed with Taubin filtering: Filters → *Smoothing, Fairing and Deformation* → *Taubin Smooth* with the following parameters: λ (lambda) = 0.6, μ (mu) = -0.53 and 100 iterations. Paraview is used for 3D rendering.

**nnU-Net**

To automatize LSEC segmentation, we selected the Python package "nnU-Net" [1], which leverages deep convolutional neural networks (CNN). We used the reference segmentation from ilastik as "Ground Truth" to train the CNN segmentation model from nnU-Net.

The nnU-Net package requires specific steps to process the data: Setting up a dedicated Python environment with all the required dependencies (including the machine-learning library "PyTorch"); creating and referencing with environment variables the raw data, pre-processed data and trained model weights directories; and transforming arbitrary raw data into the required nnU-Net input data format. The structure of the said data format is based on the format used for the image analysis challenge from the Medical Segmentation Decathlon [2].

nnU-Net automatically scales the underlying CNN model depending on the data shape, and uses cross-validation (CV) as an unbiased technique to fine-tune the model hyperparameters during training. In particular, nnU-Net uses the so-called five-fold cross-validation, that is depicted in **Figure S5**. It divides the training data into 5 equally-sized subsets (called "folds"), and repeats the training of the model also five times. Each time, a different fold is excluded from the training data, and it is used to evaluate the segmentation performance of the model.

The package nnU-Net offers support for various CNN architectures out-of-the-box, including the famous U-Net [3]: an encoding path that progressively extracts larger scale features from the image, and a symmetric decoding path that reconstructs the segmentation by combining spatial details from various scales. While nnU-Net can automatically tune most of its parameters, some still need to be adjusted according to the characteristics of the input data. This includes the choice of various types of models (e.g., standard U-Net vs residual encoder U-nets, etc) and their dimensionality (i.e., 2D vs 3D). When working with large volumes, it might also be advisable to divide them into smaller sub-volumes. The sub-volume size requires additional careful evaluation. We use a JSON (JavaScript Object Notation) file to store all these different attributes and parameters.

The key parameters optimized according to the characteristics of our data are detailed below.

**Dimensionality**: The selection of the U-Net architecture dimensionality (i.e., 2D vs 3D) depends on the following criteria and constraints: the complexity of the target structure; the dimensionality of the data (image vs volumes), and the available computational resources (small local workstation vs large node on a HPC cluster). In our case, for instance, we worked with 3D data on a moderately old Linux workstation, equipped with a NVIDIA RTX 5000 GPU (equivalent to a NVIDIA RTX 2080 Ti). This configuration made us first consider training a 2D U-Net (using 2D convolutions), which is faster (by approx. a factor 2) and uses less GPU memory than a 3D U-Net. We used this 2D U-Net to fine-tune sub-volume sizes (i.e., patch size) and other parameters. Following this initial parameter-tuning step, we also trained a 3D U-Net, which is better suited for the complexity of the features in our data.

**Epochs**: The number of epochs is chosen to minimize the training time in order to prevent overfitting. For our data, we have observed that a 2D U-Net converges in 150 epochs (**Figure S6**), while a 3D U-net converges in 350 epochs. We have modified the nnU-Net code to allow manually choosing the number of epochs.

**Patch size**: Larger patches (sub-volumes) are preferred as they preserve more spatial context, which is crucial for accurate segmentation. However, since some of the sub-volumes are dedicated to the test and cross-validation sets, we need a sufficient number of sub-volumes in order to have enough representation for different regions of the raw volume. Thus, the user should find an equilibrium between larger and smaller patch sizes that satisfies both constraints. As mentioned above, the sub-volume split creates boundary discontinuities between patches, which can lead to fragmented segmentation. For this reason, we allow some overlap between adjacent sub-volumes, by extending them. The overlap was: (50, 50, 50).

**Optimizer**: The Adam optimizer was used [4].

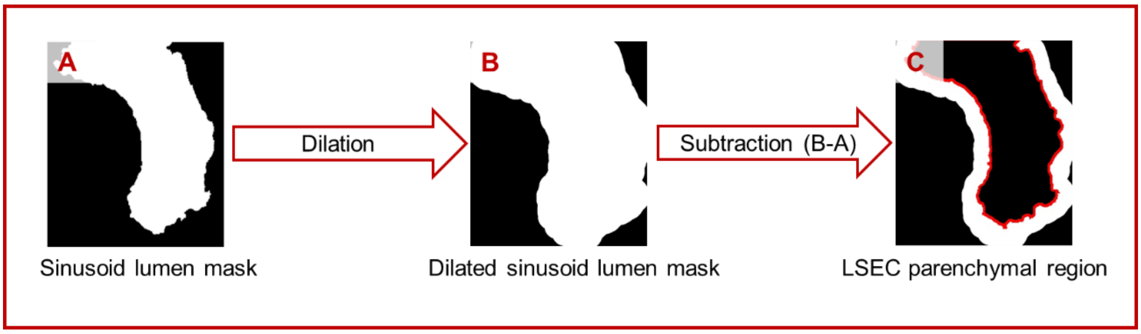


***Figure S1:*** *Strategy for the segmentation of the LSEC parenchymal region.*

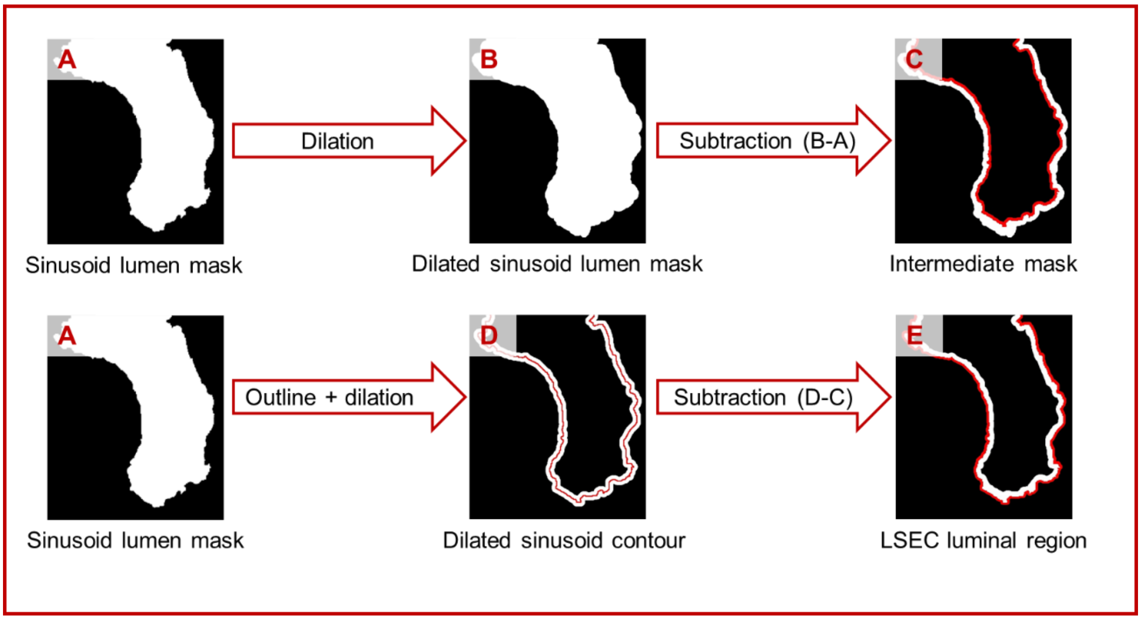


***Figure S2:*** *Strategy for the segmentation of the LSEC luminal region.*

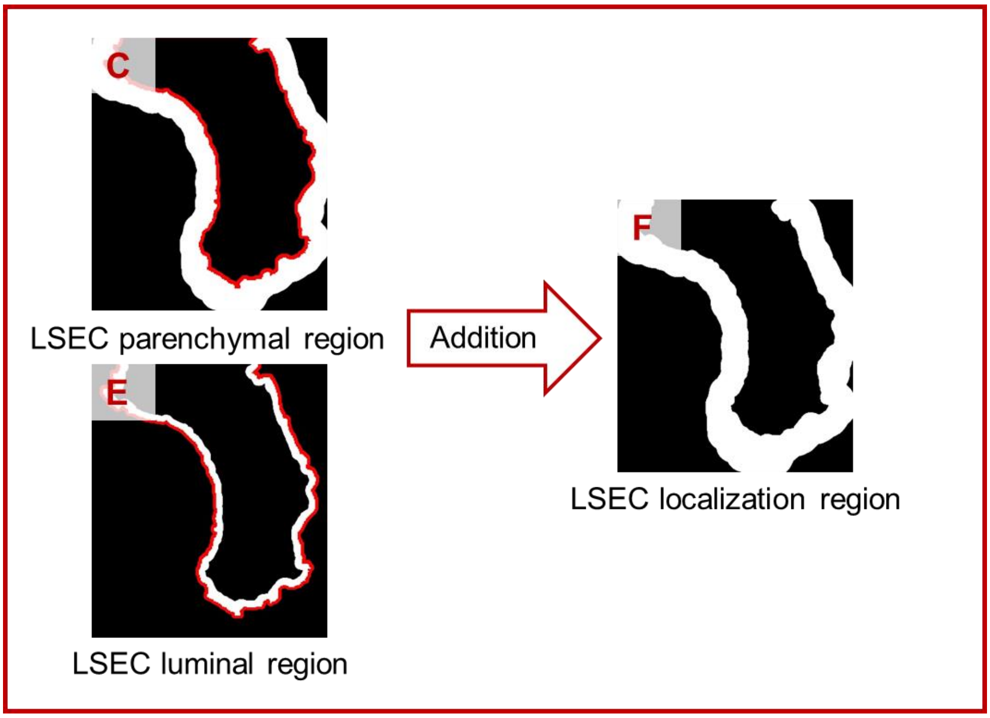


***Figure S3:*** *Strategy to determine the LSEC localization region.*

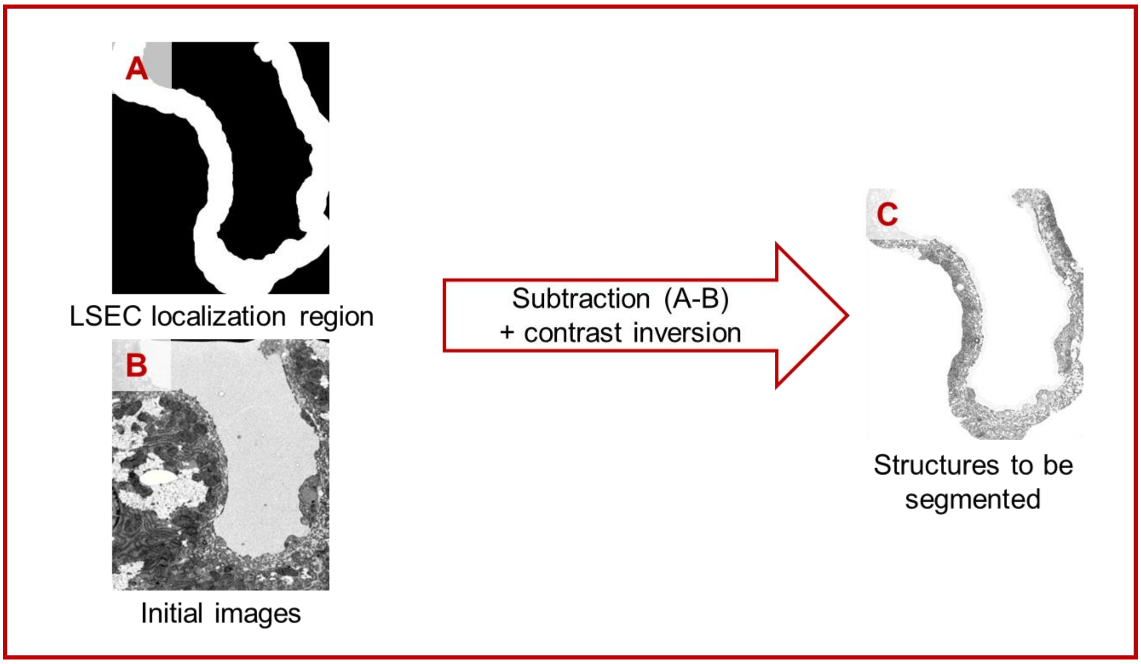


***Figure S4:*** *Extraction of the LSEC localization region in the initial data.*

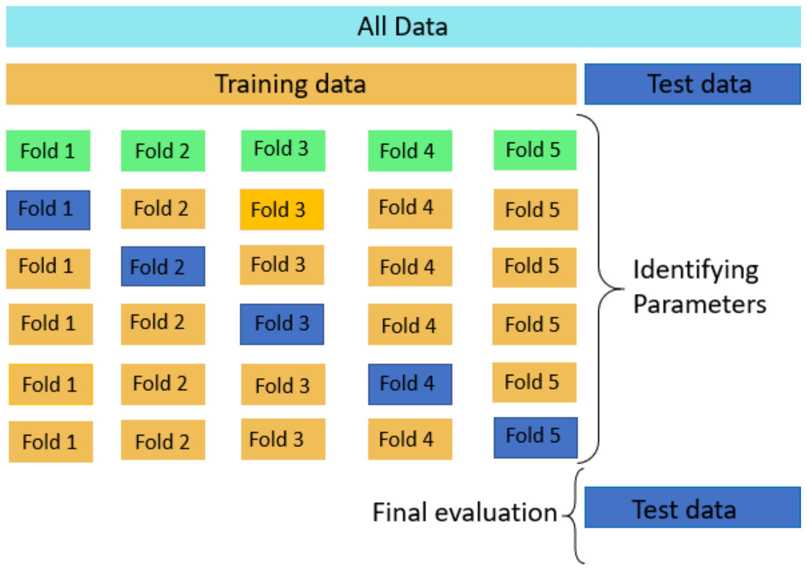


***Figure S5:*** *Training and validation using five equally sized subsets ("Folds").*

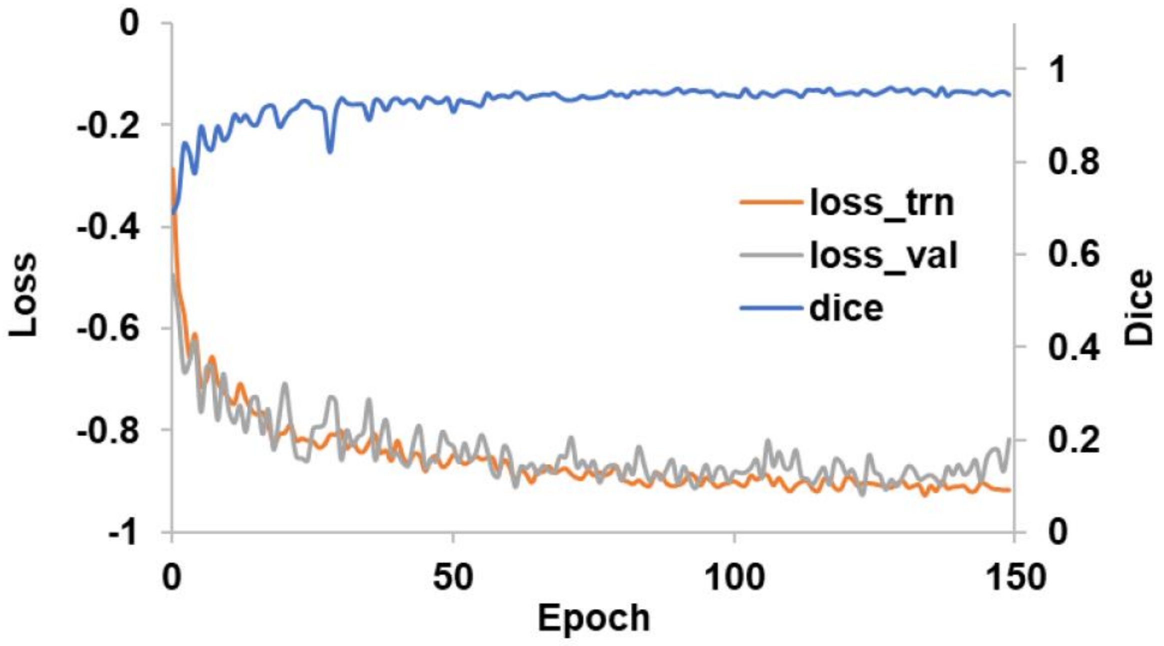


***Figure S6: Dice score and loss function values at each number of epoch during the training with a 2D U-Net configuration.*** *Under nnU-Net, the loss function is a combination: Loss = DiceLoss + CrossEntropyLoss. The Dice Loss is well suited for segmentation cases with imbalanced classes, particularly when the background predominates over the object of interest. Cross-entropy loss is used to impact classification errors at the pixel level, which helps to*

*ensure stable convergence. This score ranges from 0 to 1. The closer the score is to 1, the greater the similarity between the predictions and the ground truth. In this graph, the Dice score reaches a value of 0.98, indicating that the model is able to generalize. The higher the Dice score, the lower the loss function as shown here.*

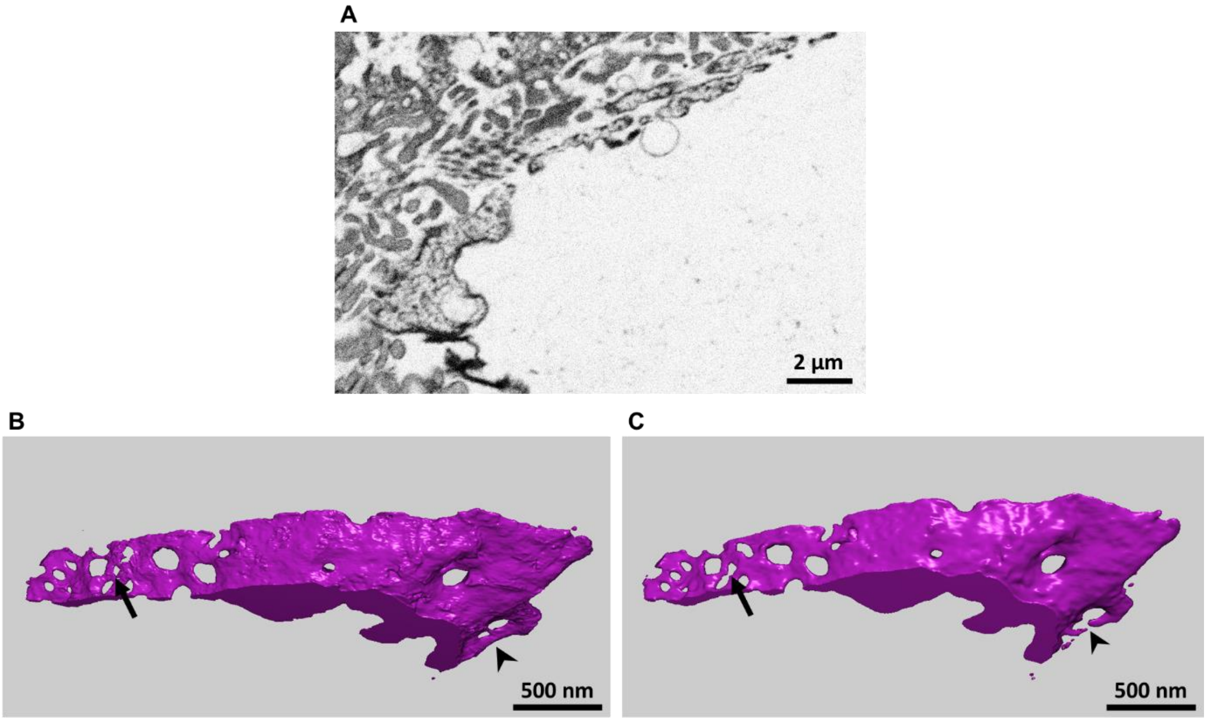


***Figure S7: Loss of LSEC fenestrae upon binning.*** *Segmentation of a region with LSEC fenestrae* ***(A)*** *using data at 4 × 4 × 4 nm³ (**B**) or binned at 8 × 8 × 8 nm³ (**C**). 3D mesh obtained for both segmentation shows fenestrae properly segmented in (**B**), while they are connected fused in (**C**).*
